# β-catenin/TCF signaling represses GUCY2C hormone expression through a p38α–GATA2 axis in colorectal cancer

**DOI:** 10.64898/2026.11.16.767112

**Authors:** Adi Caspi, Andrew E. Evans, Ariana A. Entezari, Jasmine R. Alvarez, Anne K. Londregan, Thomas J. M. Kuret, Robert D. Carlson, Jeffrey A. Rappaport, Ramkrishna Mitra, Adam E. Snook, Scott A. Waldman

## Abstract

Guanylin (GUCA2A) and uroguanylin (GUCA2B) are paracrine hormone ligands of the tumor-suppressor receptor guanylyl cyclase C (GUCY2C). In early colorectal tumorigenesis, expression of GUCA2A and GUCA2B is silenced through transcriptional repression by APC-regulated β-catenin/TCF signaling, a pathway activated early in malignant transformation. This repression depends on a locus control region (LCR) 1,000–3,683 bp upstream of *GUCA2A*. Here, we used d*<u>C</u>*as9–APEX2 *<u>b</u>*iotinylation at genomic *<u>e</u>*lements by *r*estricted *<u>s</u>*patial *t*agging (C-BERST) in colorectal cancer (CRC) cell lines with inducible suppression of β-catenin/TCF signaling to identify β-catenin/TCF-regulated proteins associated with the *GUCA2A* regulatory region that mediate repression of GUCY2C ligand transcription. Guide RNAs were used to direct dCas9-APEX2 for locus-specific proximity biotinylation of proteins. Quantitative mass spectrometry identified p38α (MAPK14) as a candidate regulator enriched near the *GUCA2A* promoter following β-catenin/TCF suppression. Loss-of-function and rescue experiments established that p38α supports GUCA2A and GUCA2B induction following β-catenin/TCF suppression. Importantly, GATA2 was identified as a p38α-dependent activator of GUCA2A and GUCA2B transcription that remained enriched near the *GUCA2A* promoter independent of β-catenin/TCF signaling. Indeed, GATA2 S192 phosphorylation was reduced by p38α loss and by active β-catenin/TCF signaling. Thus, a p38α/GATA2 axis links oncogenic β-catenin/TCF signaling to repression of GUCY2C hormone expression in CRC.

Graphical abstract
Proposed model of β-catenin/TCF-regulated repression of GUCY2C signaling. In the absence of β-catenin/TCF signaling, active p38α promotes phosphorylation of GATA2 at the *GUCA2A* promoter, upregulating GUCA2A and GUCA2B expression. Under tumorigenic conditions, β-catenin/TCF signaling is hyperactivated, suppressing the p38α/GATA2 axis and repressing transcription of GUCA2A and GUCA2B.

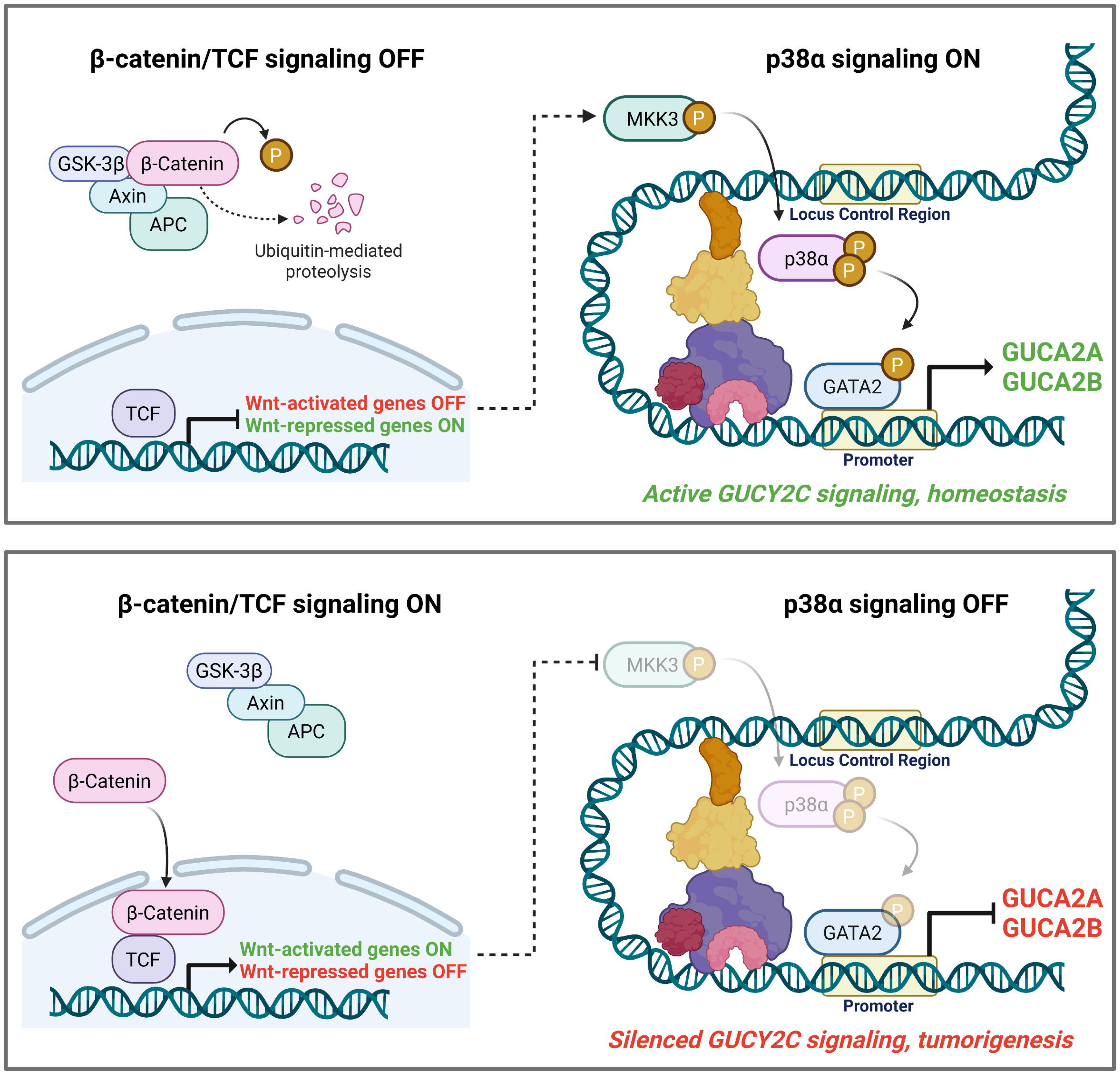

## Introduction

Colorectal cancer (CRC) is the third most commonly diagnosed cancer and the second leading cause of cancer death in the United States ^1^. In 2026, CRC was the leading cause of cancer deaths in people under 50, with a reported incidence increase of 3% annually since 2013 in the 20-49 age group ^2^. The five-year survival rate for patients with distant metastatic disease, comprising 22% of cases, is only 15% ^2^. While incidence and mortality rates have been steadily declining in older adults, the sharp rise in younger individuals and the poor prognosis associated with advanced disease underscore the need for improved prevention, detection, and early treatment options ^1^ ^2^.

APC-regulated β-catenin/TCF (β-catenin/TCF) signaling is a critical pathway driving colorectal tumorigenesis ^3^ ^4^. The β-catenin/TCF signaling cascade is a master regulator of embryonic development, tissue differentiation, and homeostasis ^4^. Aberrant activation of this pathway is a hallmark of >90% of CRC tumors, with the most frequent events being APC inactivation and β-catenin activation mutations, which promote uncontrolled cell proliferation and neoplasia ^3^ ^5^. β-catenin/TCF signaling broadly contributes to the dysregulation of apoptosis, inflammation, and cell migration through the transcriptional activation of downstream oncogenes ^3^ ^4^ ^5^. Recent studies have highlighted the repression of gene transcription by β-catenin/TCF signaling, notably through the identification of β-catenin negative regulatory elements ^6^ and a contribution from MAPK signaling ^7^. However, mechanisms through which β-catenin/TCF represses target genes in CRC remain incompletely defined.

Guanylyl cyclase C (GUCY2C) is a CRC tumor suppressor. GUCY2C, a membrane receptor of intestinal epithelial cells, regulates fluid secretion and homeostasis through cGMP signaling following activation by its ligands guanylin (GUCA2A, colon) and uroguanylin (GUCA2B, small intestine) ^8^ ^9^. GUCY2C signaling is silenced early in tumorigenesis, promoting cell-cycle dysregulation, genomic instability, inflammation, and barrier dysfunction ^10^ ^11^ ^12^ ^13^. Silencing of GUCY2C signaling in early CRC occurs by transcriptional repression of its ligands, while GUCY2C itself is retained ^14^ ^15^. Expression of GUCA2A and GUCA2B is lost following activation of the β-catenin/TCF axis, and suppression of this pathway restores GUCY2C ligand expression ^16^ ^17^ ^18^. A locus control region (LCR) upstream of *GUCA2A* mediates the β-catenin/TCF responsiveness of both ligand genes ^18^.

Transcriptional regulators of GUCY2C ligands are poorly characterized. HNF1-α activates GUCA2A expression, but thus far has not been linked to β-catenin/TCF signaling ^19^. Recently, p38α (MAPK14) was identified as a β-catenin/TCF-repressed activator of GUCA2A and GUCA2B that requires the LCR ^20^, representing a potential target for preventive or therapeutic strategies aimed at reconstituting GUCY2C signaling and restoring intestinal homeostasis. However, the proteins associated with the GUCA2A regulatory region that mediate β-catenin/TCF-dependent repression of both ligand genes remain incompletely defined.

Proximity labelling and affinity purification can identify protein complexes and interactions at the chromatin level ^21^. Guide RNA (gRNA)-directed Cas9 targeting can capture specific DNA loci and interrogate the regulatory proteome of a DNA sequence. Strategies to “reverse” chromatin immunoprecipitation (ChIP) to capture a defined locus and identify its associated proteins without prespecifying protein targets include R-ChIP ^22^, enChIP ^23^, and CRISPR-ChAP ^24^. These techniques can be limited by factors such as antibody interaction strength, chemical crosslinking, and long reaction times, which can hinder the capture of transcription factors. Proximity labelling tools, such as biotin ligases and peroxidases, can mitigate these limitations by introducing rapid reaction kinetics and relying on strong biotin-streptavidin binding, facilitating the detection of transient or weakly associated proteins ^21^. These methods include CAPTURE ^25^, GLoPro ^26^, CAPLOCUS ^27^, and C-BERST ^28^, which have been used to map interactions with the LINE-1 element ^29^, *FOXQ1* ^30^, *LGALS3* ^31^, and *RIPK3* ^32^.

We employed C-BERST (d*<u>C</u>*as9–APEX2 *<u>b</u>*iotinylation at genomic *<u>e</u>*lements by *r*estricted *<u>s</u>*patial *t*agging) at the *GUCA2A* locus to identify associated proteins and define the mechanism of β-catenin/TCF regulation of GUCY2C hormone expression in CRC. Given the spatially restricted biotinylation radius of APEX2 (∼20 nm ^28^ ^26^), we characterized dCas9-APEX2 binding to multiple sites along the LCR. LC-MS/MS revealed p38α enrichment at the *GUCA2A* promoter in the absence of β-catenin/TCF signaling, consistent with its behavior as a β-catenin/TCF-repressed activator of GUCY2C ligands. We further identified GATA2 as a transcription factor enriched at the *GUCA2A* promoter whose activity is suppressed by β-catenin/TCF signaling and that regulates GUCA2A and GUCA2B transcription. Finally, we established that GATA2-dependent hormone induction requires p38α, defining a signaling axis through which β-catenin/TCF represses GUCY2C hormone expression.

## Results

### gRNA-directed occupancy of dCas9-APEX2 at the GUCA2A locus

To investigate the regulatory machinery through which β-catenin/TCF signaling suppresses GUCA2A and GUCA2B expression, we used a system for inducible β-catenin/TCF suppression in HT29 cells expressing zinc-inducible wild-type APC (**Fig. 1A**) ^33^. After 24 h of zinc treatment, APC protein increased, accompanied by β-catenin degradation and induction of GUCA2A and GUCA2B expression (**Fig. 1B-C, Supplemental Fig. 1A**) ^16^ ^18^ ^20^. We next created a stable population of cells expressing dCas9-APEX2 ^28^. Expression of dCas9-APEX2 did not interfere with zinc-dependent induction of GUCA2A or wild-type APC (**Fig. 1D**). Next, cells were transduced with constructs encoding single guide RNAs to direct dCas9-APEX2. gRNAs were designed to target candidate regulatory regions upstream of the *GUCA2A* gene, including the proximal promoter and DNase hypersensitive sites within the locus control region (LCR), approximately 1.2 kb and 3.5 kb upstream of the transcription start site (**Fig. 1E**)^18^.

**Figure 1:**
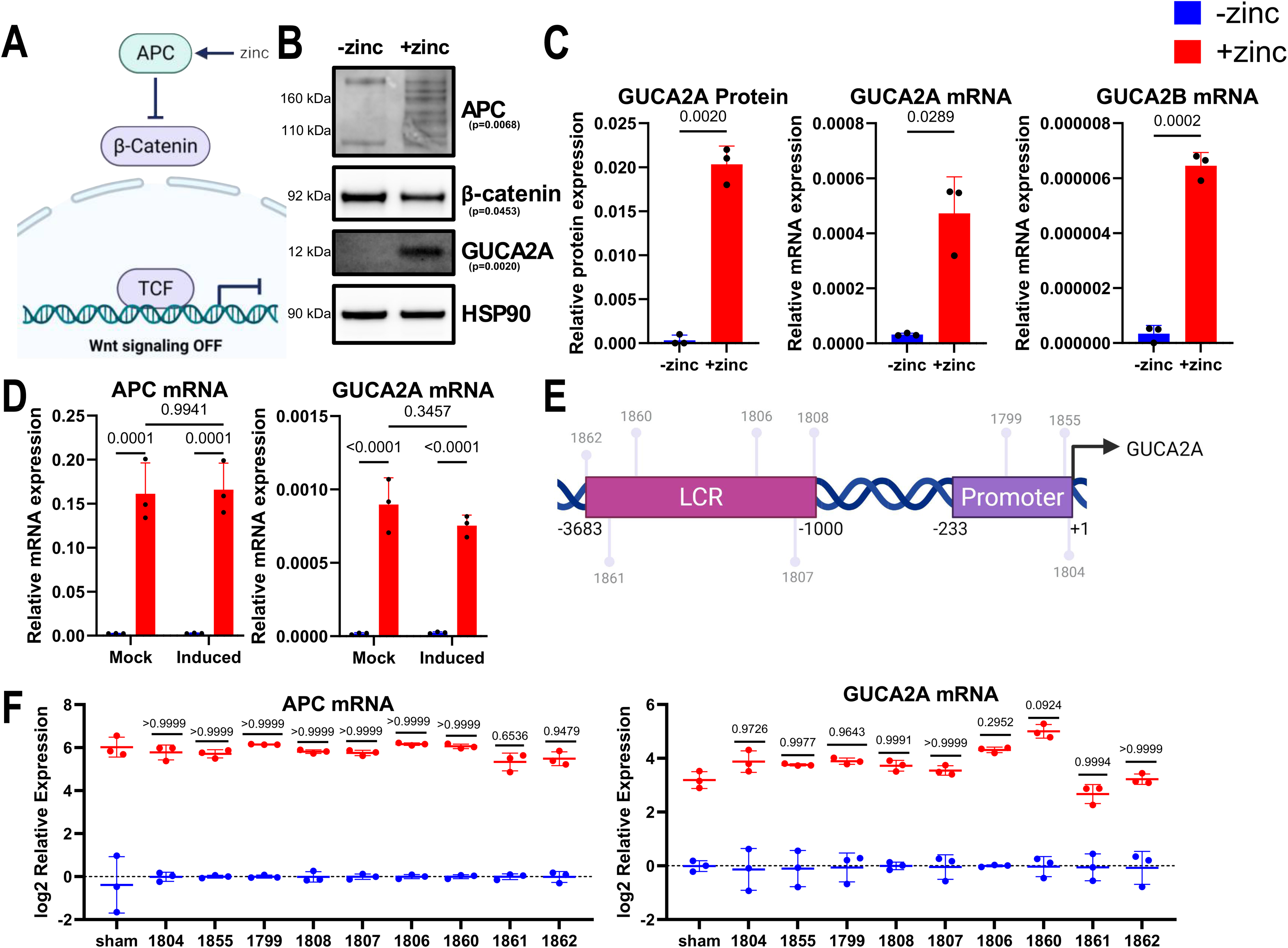
GUCA2A-targeting gRNAs preferentially recruit dCas9-APEX2 to the intended locus. (A) Schematic of HT29 (WT APC) cells. Following 24 hours of zinc treatment, APC is induced, promoting the degradation of β-catenin and suppression of downstream β-catenin/TCF signaling. (B) Immunoblot of APC induction, and downstream β-catenin degradation, in untreated and zinc-treated HT29 (WT APC) cells. Welch’s t-test p values are shown. (C) GUCA2A protein and mRNA, and GUCA2B mRNA, are induced following zinc induction of APC. (D) mRNA induction of APC (left) and GUCA2A (right) in HT29 (WT APC) cells is preserved upon doxycycline- and Shield1-dependent induction of dCas9-APEX2. (E) Layout of gRNAs along the *GUCA2A* regulatory elements: the proximal promoter (gRNAs 1799, 1855, 1804); the −1.2 kb site in the LCR (gRNAs 1806, 1807, 1808); and the −3.5 kb site in the LCR (gRNAs 1862, 1861, 1860). (F) mRNA induction of APC (left) and GUCA2A (right) is unchanged in the presence of GUCA2A-targeting gRNAs relative to a non-targeting gRNA (sham). For each gRNA, relative expression is shown as the zinc condition log2-fold normalized to its respective no-zinc condition. p values compare the zinc condition for each gRNA to the zinc condition for the sham gRNA. Welch’s t-test (C); two-way ANOVA with Tukey’s correction for multiple comparisons (D, F). N=3 biological replicates per condition (B-D, F) with 2 technical replicates per sample in qPCR (C, D, F).

We assessed whether targeted gRNAs preserved zinc-dependent induction of GUCA2A expression. Given that dCas9 can sterically interfere with trans-acting factors ^34^, we sought to ensure that gRNAs driving dCas9-APEX2 to the *GUCA2A* regulatory region preserved hormone induction as a functional check for interference with critical regulators. Nine *GUCA2A*-targeting gRNAs were identified that preserved zinc-dependent induction of APC and GUCA2A relative to a nontargeting sham gRNA (**Fig. 1F**). These results support the compatibility of C-BERST at this locus, as GUCA2A expression and β-catenin/TCF-regulated dynamics were preserved.

### Targeted gRNAs preferentially recruit dCas9-APEX2 to the GUCA2A locus

As *GUCA2A*-targeting gRNAs had not been previously validated, we characterized their ability to promote dCas9-APEX2 capture of the *GUCA2A* locus. Three gRNAs were selected to target the promoter and the −1.2 kb and −3.5 kb hypersensitivity sites of the enhancer. We performed ChIP-qPCR against the mCherry tag on dCas9-APEX2 to measure enrichment of each targeted region with its corresponding gRNA. Compared with a sham gRNA, *GUCA2A*-targeting gRNAs demonstrated strong enrichment at their targeted sites along the *GUCA2A* regulatory region (**Fig. 2A**). ChIP-seq corroborated these findings (**Fig. 2B**), with the center of each peak within 30 bases of the predicted gRNA binding site. To assess gRNA specificity, differential binding analysis was performed to characterize binding peaks enriched by each targeted gRNA, compared with sham. For each gRNA, the predicted *GUCA2A* site was the most significantly enriched peak, with potential off-target sites exhibiting weaker enrichment (**Fig. 2C**). Thus, gRNAs designed against the *GUCA2A* promoter and enhancer preferentially recruit dCas9-APEX2 to their intended loci, although lower-level off-target occupancy is detectable.

**Figure 2:**
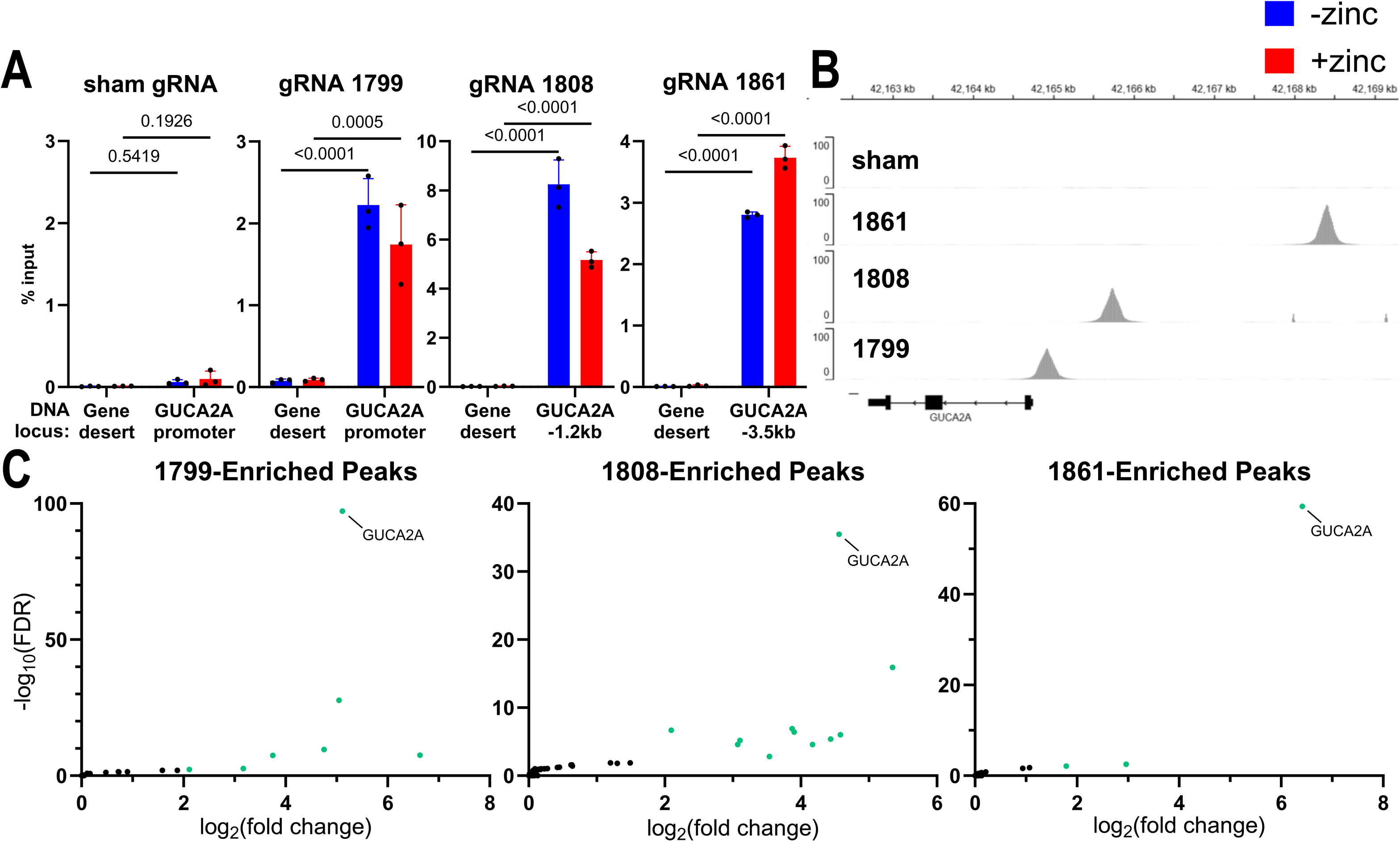
dCas9-APEX2 occupancy at the *GUCA2A* locus enables effective and relatively specific targeted DNA capture. (A) Anti-mCherry ChIP-qPCR of dCas9-APEX2 in HT29 (WT APC) cells with gRNAs against the *GUCA2A* promoter (1799), proximal LCR (1808), or distal LCR (1861) compared to sham. For each targeted gRNA, pulldown is probed by primers flanking its respective targeted site. A gene desert on chromosome 12 serves as a negative control. Two-way ANOVA with Tukey’s correction for multiple comparisons. (B) Snapshot of anti-mCherry ChIP-seq peaks at the *GUCA2A* locus using reads per million (RPM)-normalized tracks. (C) Differential binding analysis of ChIP-seq peaks enriched by each targeted gRNA relative to sham. Significantly enriched peaks (green) met thresholds of fold change (FC)>2 and false discovery rate (FDR)<0.01. The predicted *GUCA2A* peak for each gRNA is denoted. N=3 replicate IPs per condition with 2 technical replicates per sample in qPCR (A); N=2 replicate IPs per condition (B-C).

### C-BERST at the GUCA2A locus

We next performed C-BERST (**Fig. 3A**) ^28^, using promoter-targeted dCas9-APEX2 to biotinylate proximal proteins following biotin phenol (BP) and H_2_O_2_ treatment and quenching. Immunoblot analysis of whole-cell lysates revealed abundant biotinylation of the proteome (**Fig. 3B**). Notably, biotinylation was also abundant in sham gRNA-expressing cells, consistent with previous reports of APEX2-mediated background labelling ^28^ ^26^ ^35^ ^36^. Biotinylation was markedly reduced in the absence of exogenous BP, with strong bands corresponding to known endogenously biotinylated proteins ^37^. By immunofluorescence, streptavidin signal was confined to the nucleus and colocalized with dCas9-APEX2, while cells not treated with BP lacked detectable streptavidin signal (**Fig. 3C**).

**Figure 3:**
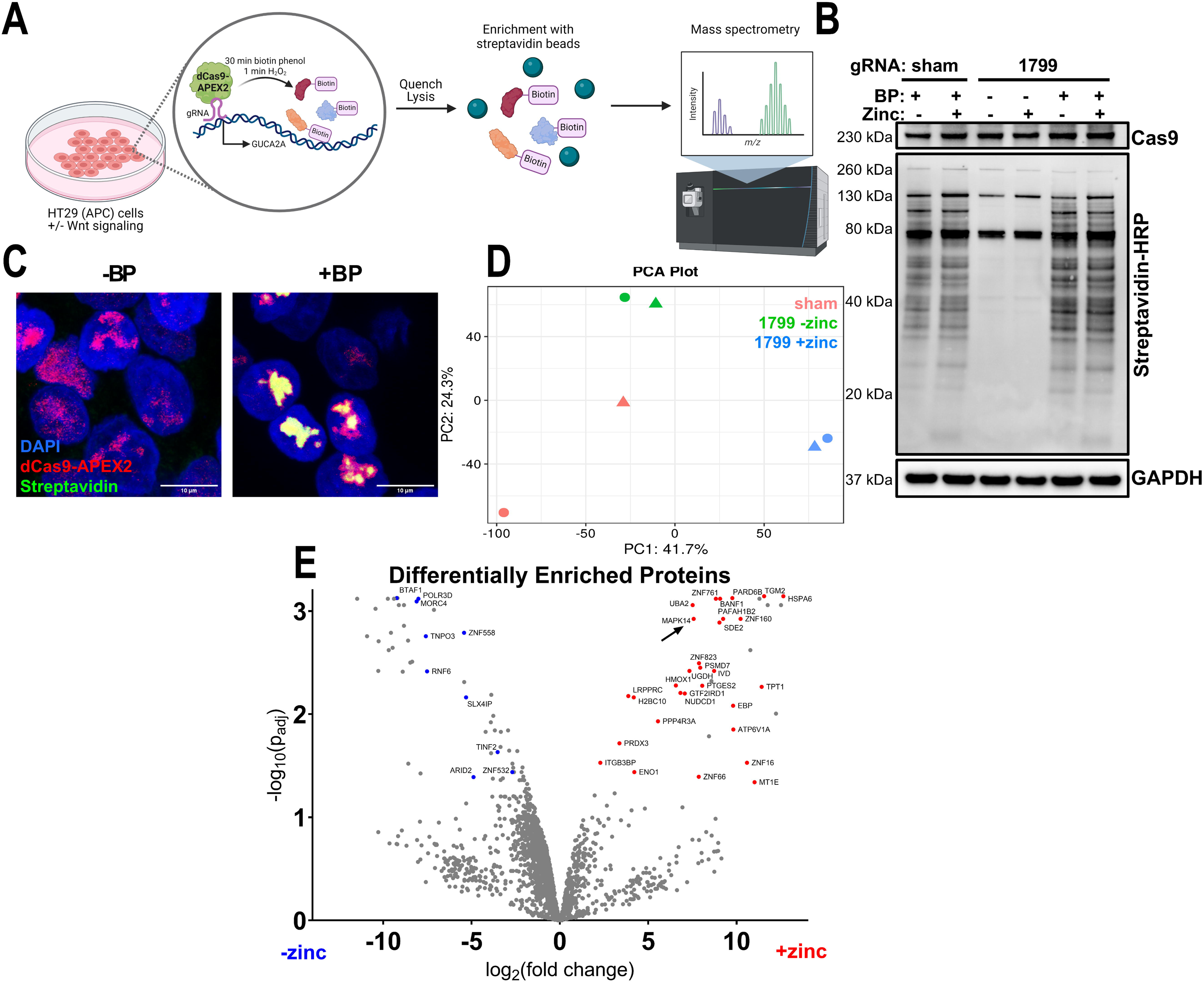
C-BERST reveals candidate mediators of β-catenin/TCF-regulated GUCA2A expression. (A) Schematic of C-BERST. HT29 (WT APC) cells expressing dCas9-APEX2 and a gRNA are treated with BP, then with H_2_O_2_ to induce proximity biotinylation. After quenching and lysis, biotinylated proteins are enriched with a streptavidin pulldown and evaluated by LC-MS/MS. (B) Immunoblot of biotinylated proteins in whole-cell lysates following C-BERST. In cells receiving no exogenous BP, streptavidin bands correspond to endogenously biotinylated proteins: acetyl-CoA carboxylase (∼265 kDa); pyruvate carboxylase (∼130 kDa); 3-methylcrotonyl-CoA carboxylase (∼75 kDa); and propionyl-CoA carboxylase (∼72 kDa) ^37^. (C) Immunofluorescence staining of HT29 (WT APC) cells expressing dCas9-APEX2 (mCherry) and *GUCA2A* promoter gRNA (1799) following C-BERST. Left: no exogenous BP; right: with BP treatment. (D) Principal component analysis of LC-MS/MS data following C-BERST with a sham or *GUCA2A* promoter gRNA (1799). (E) Enrichment of proteins with a *GUCA2A* promoter gRNA (1799). Denoted candidates (blue, red) met thresholds of fold change abundance in the experimental conditions of >2, p_adj_<0.05, and fold change abundance >2 over sham, i.e., non-background. N=2 replicates per condition in C-BERST proteomics (D-E).

C-BERST followed by streptavidin enrichment and proteomics was performed in cells expressing either sham gRNA or *GUCA2A* promoter-targeting gRNA with or without zinc induction. Principal component analysis of LC-MS/MS data demonstrated clustering of replicate experimental samples and divergence based on zinc status (**Fig. 3D**), consistent with changes in the captured proteome following β-catenin/TCF suppression. Sham replicates were more dispersed, indicating variability in the background captured proteome (**Fig. 3D**).

We analyzed C-BERST proteomic data with a focus on proteins differentially enriched at the *GUCA2A* promoter with or without zinc to reveal candidate β-catenin/TCF-dependent regulators of expression (**Fig. 3E**). Candidates enriched in the presence of β-catenin/TCF signaling (without zinc induction) were considered candidate repressors, potentially involved in the silencing of the GUCY2C hormones in tumorigenesis. These included the transcriptional repressor and chromatin remodeler BTAF1 ^38^; the tumor-promoting, DNA-binding protein MORC4 ^39^; and the β-catenin/TCF-associated oncogenic regulator RNF6 ^40^. Conversely, proteins enriched in the absence of β-catenin/TCF signaling (with zinc) were considered candidate activators that might support GUCA2A and GUCA2B expression following β-catenin/TCF suppression. These included the tumor suppressors HSPA6 and SMEK1 (PPP4R3A) ^41^ ^42^; nuclear proteins with context-dependent roles, such as SDE2 ^43^ ^44^ and ZNF16 ^45^; as well as other transcriptional modulators, including TPT1 ^46^ and GTF2IRD1 ^47^.

### p38α is a β-catenin/TCF-repressed activator of GUCA2A and GUCA2B

p38α (MAPK14) was differentially enriched at the *GUCA2A* promoter in the absence of β-catenin/TCF signaling (**Fig. 3E**). p38α, a member of the MAPK family, has been implicated in regulating proliferation and differentiation of the intestinal epithelium, as well as the development of CRC and inflammatory bowel disease ^48^ ^49^ ^50^ ^51^ ^52^. In early tumorigenesis, p38α acts as a tumor suppressor through its capacity to regulate apoptosis and the cell cycle ^51^ ^52^. Nuclear p38α complexes with transcription factors at target gene loci, modulating their activity through phosphorylation ^48^ ^53^. Recently, we reported that p38α activity was suppressed by β-catenin/TCF signaling in CRC, and active p38α upregulates GUCA2A and GUCA2B expression ^20^. Previously, p38α protein was detected at the *GUCA2A* locus by CAPTURE immunoblot, another dCas9 affinity capture technique ^20^ ^25^. Identification of p38α at the *GUCA2A* promoter by C-BERST supports these observations, reinforcing a role for p38α in β-catenin/TCF regulation of GUCA2A and GUCA2B expression.

To validate these observations, and to address potential false-positive findings from off-target binding sites (**Fig. 2C**), we performed siRNA knockdown of C-BERST candidate regulators. For high-throughput screening, we used a more readily transfectable model of inducible β-catenin/TCF suppression, LS174T cells expressing a doxycycline (DOX)-inducible dominant negative TCF (DNTCF) ^54^. Like HT29 cells with zinc-inducible APC, these cells induce GUCA2A and GUCA2B expression following DOX treatment (**Supplemental Fig. 1B-D**) ^16^ ^18^ ^20^. Two independent siRNAs were used per target to assess concordance of target-specific effects. The siRNA screen was combined with a GUCA2A reporter construct in which the 6-kb upstream regulatory region of *GUCA2A*, including the LCR, drives luciferase expression ^18^. siRNA knockdown of p38α reduced the DOX-driven induction of luciferase activity (**Fig. 4A; Supplemental Fig. 2A**). In contrast, other candidates did not meet the prespecified criterion of concordant effects with both siRNAs (**Supplemental Fig. 2A**). Knockdown of p38α also reduced DOX induction of endogenous GUCA2A and GUCA2B (**Fig. 4B; Supplemental Fig. 2B**), reinforcing the finding that p38α is required for full induction of GUCY2C hormone expression following β-catenin/TCF suppression.

**Figure 4:**
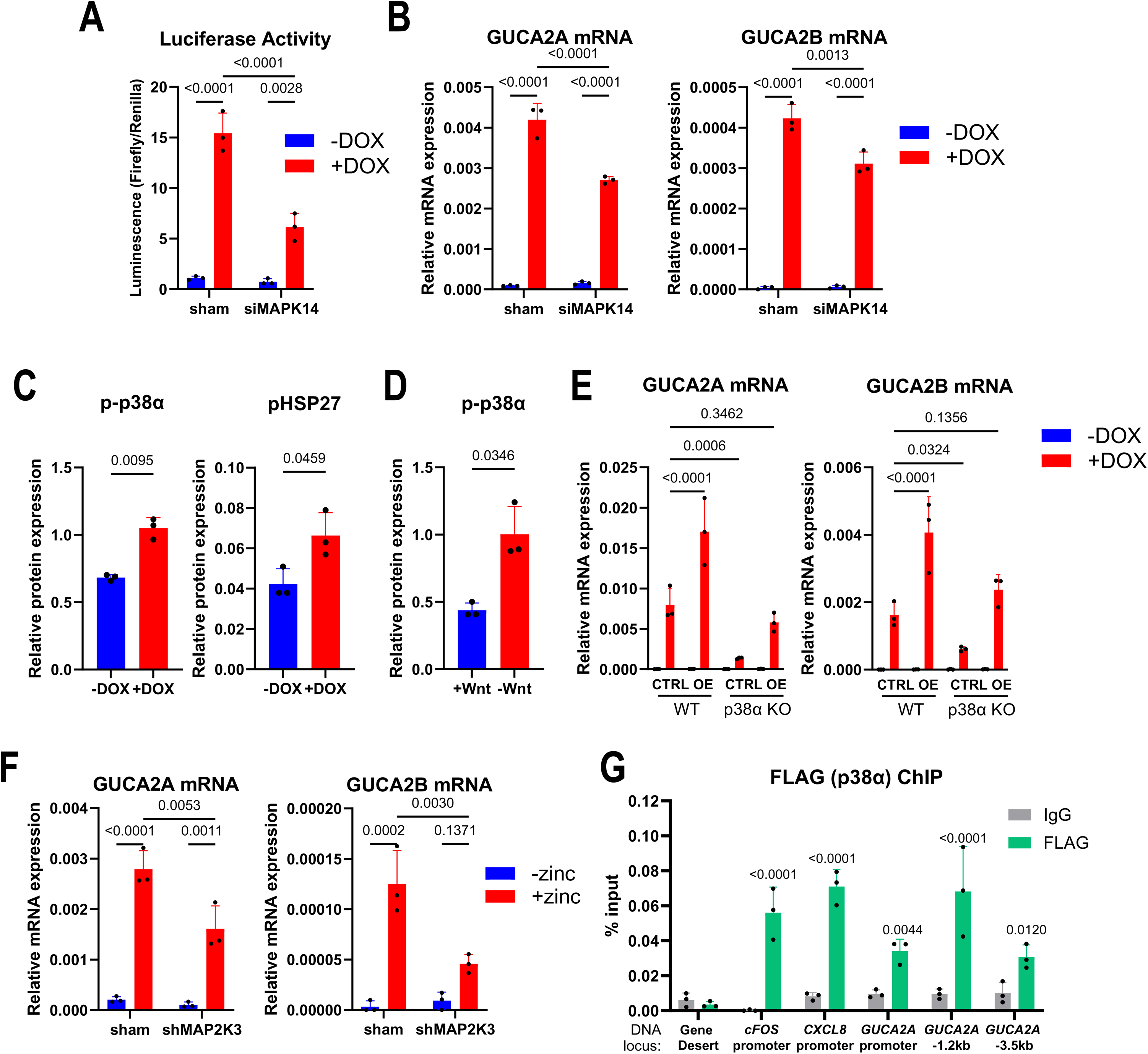
p38α is a β-catenin/TCF-repressed activator of GUCA2A and GUCA2B expression. (A) Luciferase activity of a 6-kb GUCA2A reporter vector in LS174T (DNTCF) cells is reduced in the presence of p38α-targeted siRNA. (B) GUCA2A and GUCA2B mRNA induction is reduced in LS174T (DNTCF) cells in the presence of p38α-targeted siRNA. (C) Phosphorylated p38α (T180/Y182) and downstream HSP27 (S82) are suppressed by β-catenin/TCF signaling in LS174T (DNTCF) cells. (D) p38α phosphorylation is reduced under proliferative conditions in colonic organoids. (E) GUCA2A and GUCA2B mRNA in control or p38α-overexpressing (OE) wild-type and p38α KO LS174T (DNTCF) cells. (F) GUCA2A and GUCA2B mRNA induction is reduced in the presence of MAP2K3-targeted shRNA in HT29 (WT APC) cells. (G) Anti-FLAG ChIP-qPCR in FLAG-p38α-expressing, p38α KO LS174T (DNTCF) cells. Anti-IgG ChIP, and a gene desert on chromosome 12, serve as negative controls. Denoted p values compare each pulldown to anti-FLAG pulldown of the gene desert. Two-way ANOVA with Tukey’s correction for multiple comparisons (A-B, E-G); Welch’s t-test (C-D). N=3 biological replicates per condition (A-G) with 2 technical replicates per sample in qPCR (B, E-G).

The β-catenin/TCF-dependent presence of p38α near the *GUCA2A* promoter (Fig. 3E) suggests that β-catenin/TCF signaling limits its association with this region. Baseline p38α expression was unaffected by β-catenin/TCF signaling; however, p38α activity, assessed by phosphorylation at T180/Y182 ^55^, and phosphorylation of the downstream pathway readout HSP27 ^56^, increased following β-catenin/TCF suppression (**Fig. 4C; Supplemental Fig. 1A, 1C, 2C**). Similarly, p38α phosphorylation was higher in differentiated mouse colonic organoids cultured without Wnt than in proliferating organoids cultured with Wnt (**Fig. 4D; Supplemental Fig. 2D**). These findings support the model that β-catenin/TCF signaling suppresses p38α activity, contributing to GUCY2C hormone loss in tumorigenesis.

To determine whether restoring p38α rescues hormone expression, we used p38α CRISPR knockout (KO) cells ^20^ and overexpressed p38α. CRISPR KO of p38α reduced the DOX-driven induction of GUCA2A and GUCA2B (**Fig. 4E, Supplemental Fig. 2E-F**), recapitulating knockdown of p38α. In both wild-type and KO cells, overexpression of p38α increased DOX induction of GUCA2A and GUCA2B, with expression in rescued KO cells comparable to that in control wild-type cells under β-catenin/TCF-off conditions (**Fig. 4E**). However, p38α overexpression did not rescue GUCA2A or GUCA2B suppressed by β-catenin/TCF signaling, suggesting that p38α is required, but not sufficient, to restore transcription under active β-catenin/TCF signaling.

To further investigate the mechanism of p38α repression by β-catenin/TCF signaling, we explored kinases upstream of p38α. p38α is canonically activated by MKK3 (MAP2K3), MKK4 (MAP2K4), and MKK6 (MAP2K6) ^48^ ^49^ ^57^. As MKK6 is expressed at low levels in HT29 (WT APC) cells (**Supplemental Fig. 3A**), we focused on MKK3 and MKK4. Like p38α, phosphorylation of MKK3 and MKK4 was suppressed by β-catenin/TCF signaling (**Supplemental Fig. 3B**). shRNA knockdown of MKK3 phenocopied loss of p38α and reduced GUCA2A and GUCA2B expression, supporting a role for MKK3 in hormone induction in the absence of β-catenin/TCF signaling (**Fig. 4F, Supplemental Fig. 3C**). In contrast, MKK4 knockdown increased the induction of GUCA2A (**Supplemental Fig. 3D**). As MKK4 is reported to require a different mechanism of activation than MKK3/6 ^57^, this finding raises the possibility of distinct MKK4-dependent regulation of hormone expression in this CRC model.

We sought to map the association between p38α and the *GUCA2A* locus using chromatin immunoprecipitation (ChIP). We used FLAG-p38α-expressing p38α KO cells to perform anti-FLAG ChIP and probed for pulldown along the *GUCA2A* regulatory element (**Fig. 4G, Supplemental Fig. 2E**). The *FOS* and *CXCL8* promoters were used as positive control loci for p38α association ^53^. Consistent with C-BERST, p38α was identified at the *GUCA2A* locus, with the strongest enrichment detected at the −1.2 kb site (**Fig. 4G**). Given that C-BERST was performed with a *GUCA2A* promoter-targeting gRNA, this finding raises the possibility of topological proximity between the promoter and enhancer, which remains to be tested directly. Overall, these observations identify p38α as a β-catenin/TCF-repressed regulator required for full GUCA2A and GUCA2B induction.

### GATA2 is a β-catenin/TCF-repressed activator of GUCA2A and GUCA2B

Given that p38α is a kinase that regulates transcription factor activity, we sought to identify transcription factors from the C-BERST data that might operate downstream of p38α. The siRNA screen did not reveal additional candidates with the ability to independently regulate *GUCA2A* reporter activity upon knockdown (**Supplemental Fig. 2A**). We analyzed the proteomic data to identify additional candidates whose enrichment at the promoter is not dependent on β-catenin/TCF signaling. While some p38α targets, such as c-Fos and JunB, require p38α for their recruitment to DNA, others, such as CHOP and MRF4, do not ^58^ ^59^ ^60^. Therefore, we hypothesized that transcription factors may remain associated with the GUCA2A promoter irrespective of β-catenin/TCF signaling, and require activation by p38α to modulate GUCA2A and GUCA2B transcription. Such factors could be missed in the differential enrichment analysis; thus, we modified the analysis to reveal all proteins enriched at the promoter over sham in either experimental condition (**Fig. 5A**). Previously identified candidates, which were differentially enriched, populated the plot for their respective enriched condition (e.g., p38α (MAPK14) is identified only in **Fig. 5A**, right). However, other proteins were enriched over sham in both datasets, consistent with promoter-proximal association under both β-catenin/TCF conditions. These included the transcriptional regulators SAP30BP, PAXIP1, ZNF275, and DNTTIP1 ^61^ ^62^ ^63^ ^64^.

**Figure 5:**
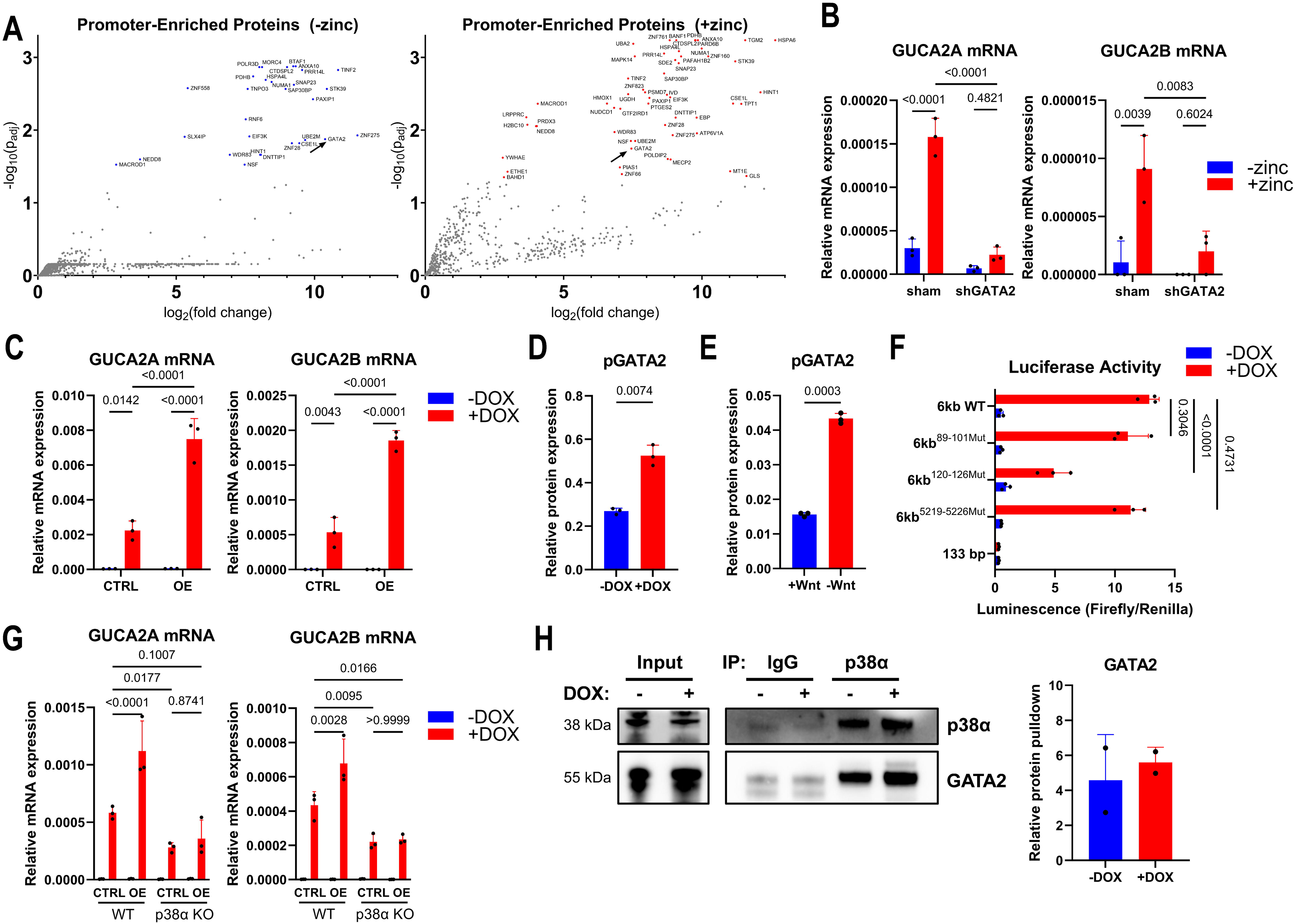
GATA2 is a β-catenin/TCF-repressed activator of GUCA2A and GUCA2B downstream of p38α. (A) Enrichment of proteins with a *GUCA2A* promoter gRNA (1799) over sham without zinc (left) or with zinc (right) by C-BERST in HT29 (WT APC) cells. Denoted candidates met thresholds of fold change abundance of >2 and p_adj_<0.05. (B) GUCA2A and GUCA2B mRNA induction is reduced by GATA2 shRNA in HT29 (WT APC) cells. (C) GUCA2A and GUCA2B mRNA induction is increased in GATA2-overexpressing (OE) LS174T (DNTCF) cells. (D) Phosphorylated GATA2 (S192) is suppressed by β-catenin/TCF signaling in LS174T (DNTCF) cells expressing FLAG-GATA2. (E) GATA2 phosphorylation is lower under proliferative than differentiated conditions in colonic organoids. (F) Luciferase activity in LS174T (DNTCF) cells of a wild-type (WT) 6-kb *GUCA2A* reporter vector and with mutant GATA2 consensus binding sites at 89-101, 120-126, and 5219-5226 bases upstream of the transcription start site. A β-catenin/TCF-unresponsive vector consisting of the *GUCA2A* proximal promoter (133 bases) driving luciferase expression is used as a negative control. (G) GUCA2A and GUCA2B mRNA induction is increased by GATA2 overexpression (OE) in wild-type LS174T (DNTCF) cells, but not in p38α KO cells. (H) Co-immunoprecipitation with an isotype control or p38α antibody in LS174T (DNTCF) cells overexpressing GATA2. Quantification shows pulldown of GATA2 relative to pulldown of p38α in either condition. Two-way ANOVA with Tukey’s correction for multiple comparisons (B-C, F-G); Welch’s t-test (D-E). N=2 replicate samples per condition (A, H). N=3 biological replicates per condition (B-G) with 2 technical replicates per sample in qPCR (B-C, G).

Among these, GATA2 emerged as a transcription factor potentially involved in β-catenin/TCF regulation of GUCA2A and GUCA2B. GATA2 is a member of the GATA family of transcription factors, a highly conserved group of proteins involved in development and tissue differentiation ^65^. GATA2 is primarily characterized as a regulator of hematopoiesis, but also has been implicated in lung, prostate, endometrial, and pancreatic cancers, with opposing roles reported in the literature ^66^ ^67^ ^68^ ^69^. ChIP-seq datasets in the ChIP-Atlas ^70^ from various cell lines identified GATA2 binding at the *GUCA2A* locus (**Supplemental Fig. 4A**). Four instances of the GATA2 consensus site, (A/T)GATAAG ^65^ ^71^, were identified by FIMO motif search ^72^ within the 6-kb regulatory region upstream of *GUCA2A*, including one occurrence of an imperfect palindromic repeat in the proximal promoter (**Supplemental Fig. 4B-C**). We therefore tested whether GATA2 regulates GUCY2C hormone expression in a β-catenin/TCF-dependent manner. Knockdown of GATA2 reduced mRNA levels of GUCA2A and GUCA2B in a β-catenin/TCF-off context (**Fig. 5B, Supplemental Fig. 4D**), supporting a role for GATA2 in hormone expression. Further, treatment with the GATA inhibitor K-7174 ^73^ reduced luciferase activity of the *GUCA2A* 6-kb reporter, along with GUCA2A and GUCA2B mRNA (**Supplemental Fig. 4E**). Conversely, expression of FLAG-tagged GATA2 increased the induction of GUCA2A and GUCA2B mRNA (**Fig. 5C**). As overexpression was unable to restore hormone mRNA expression in the presence of β-catenin/TCF signaling (without DOX), these findings indicate that GATA2, like p38α, is required for full hormone induction, but increased GATA2 abundance alone is insufficient to restore ligand expression repressed by β-catenin/TCF signaling.

As β-catenin/TCF signaling did not regulate GATA2 enrichment near the *GUCA2A* promoter (**Fig. 5A**), we tested whether β-catenin/TCF signaling instead modulates GATA2 activity through posttranslational modification. While GATA2 activity is controlled by phosphorylation at residues including S73, S119, S192, and S340 ^74^ ^75^ ^76^, phosphorylation at S192 is the best characterized ^77^. In cells expressing FLAG-tagged GATA2, S192 phosphorylation increased following β-catenin/TCF suppression (**Fig. 5D, Supplemental Fig. 4F**), pointing to β-catenin/TCF-induced suppression of GATA2 activity. Similarly, GATA2 phosphorylation was lower in Wnt-containing proliferative medium than in differentiation medium in colonic organoids (**Fig. 5E, Supplemental Fig. 2D**). These findings associate active β-catenin/TCF signaling with reduced GATA2 S192 phosphorylation, providing a candidate mechanism for impaired hormone induction.

To characterize the site of GATA2 regulation in the *GUCA2A* regulatory element, we mutated the GATA2 consensus motifs in the 6-kb luciferase reporter to determine their impact on luciferase activity (**Supplemental Fig. 4B-C**). Mutation of a GATA2 motif in the promoter 120–126 bp upstream of the transcription start site produced a ∼50% loss of β-catenin/TCF responsiveness relative to the WT construct (**Fig. 5F**), implicating this site in β-catenin/TCF-responsive GUCA2A reporter activity. Together with C-BERST, this result supports a contribution of promoter-associated GATA2 to β-catenin/TCF-responsive hormone expression.

### GATA2 is regulated by, and functions downstream of, p38α

Given that p38α and GATA2 both upregulate GUCA2A and GUCA2B expression following β-catenin/TCF suppression, we sought to determine whether they act independently or are part of the same pathway. Previous studies link p38α-dependent GATA2 S192 phosphorylation to increased transcriptional activity ^74^ ^75^ ^76^. Consistent with this model, endogenous phosphorylated GATA2 was reduced in p38α KO cells relative to WT cells (**Supplemental Fig. 4G**), suggesting that p38α is needed for full GATA2 activity. Next, we sought to determine whether overexpression of GATA2 would be sufficient to rescue GUCA2A and GUCA2B expression in the absence of p38α. While GATA2 overexpression increased GUCA2A and GUCA2B induction in WT cells (**Fig. 5C**), it did not in p38α KO cells (**Fig. 5G**), supporting a requirement for p38α in GATA2-dependent hormone induction. Finally, using a co-immunoprecipitation assay, we detected an association between p38α and GATA2 (**Fig. 5H**). Direct phosphorylation of GATA2 by p38α and the requirement for S192 in hormone induction remain to be established. Nevertheless, these observations suggest a model in which GATA2 remains associated with the GUCA2A regulatory region, while p38α supports its phosphorylation and transcriptional activity. In cancer cells, APC/β-catenin/TCF signaling suppresses p38α activity, reduces GATA2 phosphorylation, and contributes to repression of GUCY2C hormone expression.

## Discussion

Activation of β-catenin/TCF signaling through APC loss or activating β-catenin mutations early in colorectal tumorigenesis initiates signaling changes that contribute to disease development ^3^ ^4^. Following APC inactivation, the GUCY2C signaling axis is silenced through loss of the colonic hormone GUCA2A and its small-intestinal counterpart GUCA2B, resulting in hyperproliferation of the crypt compartment ^10^ ^12^ ^14^ ^15^ ^16^ ^17^ ^18^. Targeted reactivation of the GUCY2C axis reversed this effect and prevented tumorigenic progression ^10^ ^11^. Thus, characterization of mechanisms underlying GUCA2A and GUCA2B loss could identify opportunities to restore this tumor-suppressive pathway. Previously, we defined the *cis*-regulatory element involved in β-catenin/TCF suppression as an enhancer (LCR) upstream of the *GUCA2A* gene locus ^18^. Further, we determined that the MAPK protein p38α regulates GUCA2A and GUCA2B expression in a β-catenin/TCF-repressed manner ^20^. Here, we validated these observations using locus-targeted proteomics (C-BERST) at the *GUCA2A* regulatory element. Indeed, p38α protein was enriched at the *GUCA2A* promoter in the absence of β-catenin/TCF signaling, consistent with a role in supporting GUCA2A and GUCA2B transcription. Upregulation of both hormones required p38α, whose activating phosphorylation was reduced by β-catenin/TCF signaling.

These observations support previous work characterizing crosstalk between β-catenin/TCF and MAPK. Indeed, the ERK and JNK arms of the MAPK signaling cascade mediate extensive transcriptional repression by β-catenin/TCF signaling in pancreatic and CRC models ^7^. Interestingly, our recent observations indicate a complementary role for the p38 branch of MAPK signaling, in which ∼75% of the β-catenin/TCF-regulated transcriptome, including GUCA2A, requires p38α signaling ^20^. Moreover, an inverse relationship between β-catenin/TCF and MAPK was reported recently in metastatic models, with a transition from a high-Wnt/low-MAPK state to a low-Wnt/high-MAPK state ^78^. This observation may help reconcile context-dependent roles for p38, which suppresses early tumorigenesis, but can support progression in established tumors ^51^ ^52^ ^48^. As tumors continue to develop, they acquire additional mutations in *KRAS*, *PIK3CA*, and *TP53*, making them less exclusively reliant on β-catenin/TCF to maintain proliferation and survival ^3^. Thus, modulation of β-catenin/TCF, p38α, and GUCY2C signaling warrants investigation in the preventive setting or in early disease.

The direct link between β-catenin/TCF and p38α in this context remains unresolved. In our models, phosphorylation of the upstream kinases MKK3 and MKK4 is reduced under active β-catenin/TCF signaling. In canonical MAPK signaling, these are activated by MAP3Ks (MAP kinase kinase kinases), which are themselves regulated by cell surface receptors ^48^. Because regulation of p38α was reproduced by manipulating TCF, the transcriptional effector of β-catenin/TCF signaling (**Fig. 4C**), we hypothesize that TCF-regulated gene products suppress the p38α axis. Future studies should determine whether upstream regulation occurs in a cell-autonomous manner or requires a receptor-mediated event to suppress p38α activity.

In contrast to p38α, GATA2 was enriched near the *GUCA2A* promoter under both β-catenin/TCF-ON and β-catenin/TCF-OFF conditions. β-catenin/TCF signaling inhibits GATA factors in cardiac muscle development ^79^. Recently, loss of GATA factors was implicated in CRC metastasis, with reduced chromatin accessibility at GATA2 consensus sites ^80^. GATA2 expression was unchanged between primary tumors and metastases; the altered motif accessibility could instead reflect changes in GATA2 activity or chromatin state. Similarly, we found that by suppressing p38α, β-catenin/TCF signaling reduces GATA2 phosphorylation and hormone induction. The binding of GATA2 to DNA is mediated by its highly conserved zinc finger motifs ^65^ ^76^. In acute myeloid leukemia cells, p38α increases DNA binding of GATA2 by phosphorylation at S192 ^74^ ^75^. Here, GATA2 enrichment near the *GUCA2A* locus persisted under both β-catenin/TCF conditions, despite differences in p38α activity. As p38α activity is not completely abrogated by β-catenin/TCF (**Fig. 4C-D, Supplemental Fig. 2C**) ^20^, baseline activity may be adequate to support GATA2 activation and DNA binding. Indeed, phosphorylated GATA2 is diminished but still present under β-catenin/TCF-ON conditions (**Fig. 5D-E**). Further, additional kinases may be responsible for phosphorylating GATA2 at S192, including those of the ERK pathway ^81^. Phosphorylation at residues besides S192 also may support DNA binding and transcriptional activity ^76^.

Overexpression of p38α or GATA2 enhanced hormone induction following β-catenin/TCF suppression but did not restore hormone expression under active β-catenin/TCF signaling. We propose that the p38α/GATA2 axis is required for full hormone induction, but additional β-catenin/TCF-dependent constraints may limit GUCA2A and GUCA2B transcription. Accordingly, increasing p38α or GATA2 abundance alone may not restore hormone expression in cells with constitutively active β-catenin/TCF signaling. Further characterization is needed to fully understand the dynamics of regulation at this locus. Chromatin architecture could contribute; for instance, both H3K27ac and H3K4me1, enhancer-associated histone marks, are strongly depleted from the *GUCA2A* locus in tumor as compared to normal tissue ^18^, consistent with an altered enhancer state that could constrain p38α/GATA2-dependent transcription. Additionally, many other transcription factors and chromatin modulators may regulate locus activity. While individual knockdown of other C-BERST candidates did not reproducibly alter reporter activity, redundant or cooperative mechanisms could mask individual contributions, warranting combinatorial perturbation studies. Indeed, many of the captured candidates include chromatin remodelers and DNA-binding proteins, which likely operate in protein complexes. Finally, beyond the promoter, proteomic targeting of elements within the LCR may reveal additional regulatory factors.

By C-BERST proteomics, both p38α and GATA2 were enriched near the *GUCA2A* proximal promoter. Consistent with this, ChIP analysis revealed p38α association at the promoter, and a GATA2 consensus motif within the promoter contributed to β-catenin/TCF sensitivity. However, β-catenin/TCF regulation of GUCA2A and GUCA2B requires the LCR, and both reporter activity and endogenous ligand expression depend on it ^18^. Interestingly, p38α also was enriched at −1.2 kb and −3.5 kb from the transcription start site, corresponding to DNase hypersensitivity elements in the LCR which regulate β-catenin/TCF-mediated repression of GUCA2A and GUCA2B ^18^. This suggests a possible promoter-enhancer contact that places LCR-associated proteins within the labeling radius of promoter-targeted APEX2 and may support hormone expression. High-resolution chromosome conformation capture techniques could test this model of chromatin architecture at this locus and the extent to which it is altered by β-catenin/TCF/p38α signaling.

These findings connect p38α-dependent GATA2 function to the loss of GUCY2C hormone transcription downstream of oncogenic β-catenin/TCF signaling in CRC. Determining how this dependency can be bypassed in β-catenin/TCF-active cells will be necessary before it can be used to restore the pathway.

## Materials and Methods

### Cell Lines

HT29 cells expressing zinc-inducible wild-type APC provided by B. Vogelstein^33^ were cultured in McCoy’s 5A (Corning #10-050-CV) supplemented with 10% FBS (Corning #35-010-CV) and 0.6 mg/mL hygromycin B (Gibco #10687010). APC induction was performed by adding 300 μM zinc chloride (Sigma-Aldrich #Z0152) for 24 hours. LS174T cells expressing doxycycline-inducible dominant negative *TCF7L2* (DNTCF) provided by H. Clevers^54^ were cultured in DMEM (Corning #10-013-CV) supplemented with 10% FBS, 10 μg/mL blasticidin S (Gibco #A1113903), and 0.5 mg/mL Zeocin selection reagent (Gibco #R25005). DNTCF was induced by adding 1 μg/mL doxycycline (DOX, Fisher Bioreagents #BP26531) for 48 hours. The GATA inhibitor K-7174 (MedChemExpress #HY-12743) was used at a concentration of 20 μM for 24 hours. Cell lines were authenticated using STR profiling (ATCC #135-XV) and tested for mycoplasma semiannually (InvivoGen #rep-mys-10).

### Plasmids

dCas9-APEX2 (pEJS578_DD-dSpyCas9-mCherry-APEX2, Addgene #108570) was packaged into lentivirus following transfection into HEK293T cells as previously described ^82^. HT29 (WT APC) cells were transduced with concentrated lentivirus and 8 μg/mL polybrene (Sigma-Aldrich #TR-1003). dCas9-APEX2 expression was induced with 2 μg/mL doxycycline and 250 nM Shield1 (Takara Bio #632189) for 21 hours. Cells were FACS sorted to select for mCherry signal (see *Fluorescence-Activated Cell Sorting*).

gRNA plasmids targeting *GUCA2A* were generated previously ^18^ and transduced into HT29 (WT APC) cells using lentiviral supernatant and 8 μg/mL polybrene. Cells expressing gRNAs were maintained in 1 μg/mL puromycin (InvivoGen #ant-pr-1) for selection. gRNA sequences are listed in **Supplemental Table 1**.

FLAG-p38α (pCDNA3 Flag p38 alpha, Addgene #20351) was transfected into LS174T (DNTCF) cells using Lipofectamine 3000 transfection reagent (Invitrogen #L3000150) according to the manufacturer’s instructions. Cells were maintained in 300 μg/mL Geneticin (Gibco #10131027) for selection.

For *GUCA2A* luciferase reporter assays, the *GUCA2A* 6-kb firefly luciferase reporter construct generated previously ^18^ was cotransfected with *Renilla* luciferase (pRL-TK, Promega #E2241) using FuGENE HD Transfection Reagent (Promega #E2311) according to the manufacturer’s instructions. A construct containing only the proximal promoter (133 bp) upstream of the luciferase gene was used as a control ^18^. Site-directed mutagenesis of the GATA2 consensus sites within the 6-kb reporter was performed using the Q5 Site-Directed Mutagenesis kit (New England Biolabs #E0554S).

pFLAG-GATA2 (Addgene #1418) was transfected into LS174T (DNTCF) cells using Lipofectamine 3000 transfection reagent according to the manufacturer’s instructions.

MISSION Lentiviral shRNAs (Sigma Aldrich #SHCLNG) were pooled and packaged into lentivirus as previously described and transduced into HT29 (WT APC) cells. Cells were maintained in 1 μg/mL puromycin for selection. shRNAs are listed in **Supplemental Table 1**.

### Fluorescence-Activated Cell Sorting (FACS)

To select for mCherry^+^ cells, cells were passed through a 70 μm filter, and single cells were counted. 1 million cells/mL were prepared for FACS with 1 μL/mL Sytox Red dead cell stain (Invitrogen #S34859). Cells were sorted on a FACSMelody Cell Sorter (BD Biosciences) with debris, doublets, and dead cells gated out. 50,000 high-mCherry^+^ cells were obtained and cultured.

### mRNA Analysis and RT-qPCR

Total RNA was isolated from cells using the RNeasy Plus Mini Kit (Qiagen #74136). 600 ng of RNA was used to generate cDNA using the TaqMan Reverse Transcription kit (Invitrogen #4304134) according to the manufacturer’s instructions. Quantitative PCR was performed using TaqMan Universal PCR Master Mix (Applied Biosystems #4318157) and TaqMan gene expression assays (primer probes listed in **Supplemental Table 1**) on a QuantStudio 3 Real-Time PCR System (Applied Biosystems #A28567) according to the manufacturer’s instructions. Quantification was performed using the delta Ct method relative to the reference gene GAPDH.

### Immunoblot Analysis

Protein was harvested from cells using M-PER buffer (Thermo Scientific #78503) supplemented with 1X Halt Protease and Phosphatase Inhibitor Cocktail (Thermo Scientific #78446). Lysates were scraped, sonicated for 10 seconds to shear DNA, and centrifuged for 10 min at 12,000g at 4°C. Protein was quantified using the Pierce BCA Protein Assay kit (Thermo Scientific #23225). 1 mg/mL lysates were prepared by diluting in NuPAGE LDS Sample Buffer (Thermo Scientific #NP0007) supplemented with β-mercaptoethanol (Gibco #21-985-023). Lysates were boiled for 10 min at 95°C and loaded onto a 4-12% NuPAGE Bis-Tris Mini Protein Gel (Invitrogen #NP0336BOX). Gels were run at 130 V for 90 min, then transferred onto a nitrocellulose transfer stack (Invitrogen #IB33001) using a 7-minute, 20 V transfer protocol on an iBlot 3 Western Blot Transfer Device (Invitrogen #A56728). Membranes were blocked for 1 hour at room temperature in 5% BSA (Fisher Scientific #BP1605-100) in TBST buffer. Membranes were incubated overnight at 4°C in primary antibodies, then for 1 hour at room temperature in secondary antibodies (listed in **Supplemental Table 1**). Membranes were exposed with SuperSignal West Dura substrate (Thermo Scientific #34076) and imaged on a ChemiDoc MP Imaging System (Bio-Rad Laboratories #12003154). Analysis was performed in Image Lab software (Bio-Rad Laboratories) by volumetric band densitometry normalized to a loading control (HSP90 or vinculin).

### Chromatin Immunoprecipitation and qPCR (ChIP-qPCR)

ChIP-qPCR was performed as previously described ^18^. Antibodies used for ChIP reactions are listed in **Supplemental Table 1**. Following DNA isolation, qPCR was run with 2 µL DNA and 200 nM forward and reverse primers (listed in **Supplemental Table 1**) using PowerUp SYBR Green Master Mix (Applied Biosystems #A25776) according to the manufacturer’s instructions on a QuantStudio 3 Real-Time PCR System. Quantification and analysis were performed using the percent input method against a 2% input sample.

### Chromatin Immunoprecipitation and Sequencing (ChIP-seq)

For ChIP-seq, cells were plated and chromatin was harvested as for ChIP-qPCR but scaled up such that each reaction consisted of lysate from a confluent 15 cm dish (20-30 million cells per IP). Chromatin was sonicated for 20 min to an average fragment length of 200-300 bp and validated on a 4150 TapeStation System (Agilent #G2992AA). For each ChIP reaction, 1200 µL chromatin supernatant was incubated with 17.6 µg mCherry polyclonal antibody (Invitrogen #PA5-34974), 2600 µL ChIP buffer (150 mM NaCl, 1% Triton X-100, 5 mM EDTA, 10 mM Tris HCl, pH 7.5), 1X Halt Protease and Phosphatase Inhibitor Cocktail, and 120 µL Dynabeads Protein G (Invitrogen #10004D). A 2% input sample from each condition was stored at -20°C. ChIP reactions were incubated overnight at 4°C on a tube rotator. The following day, the beads were washed using a MACSiMAG Separator magnetic rack (Miltenyi Biotech #130-092-168) with the following buffers: mixed micelle wash buffer, 2 times (150 mM NaCl, 20 mM Tris HCl, 5 mM EDTA, 5.2% sucrose, 1% Triton X-100, 0.2% SDS, pH 8.1); buffer 200, 2 times (0.1% w/v sodium deoxycholate, 200 mM NaCl, 0.05 M HEPES, 1 mM EDTA, 1% Triton X-100, pH 7.5); LiCl buffer, 2 times (0.5% w/v sodium deoxycholate, 250 mM LiCl, 1 mM EDTA, 0.5% v/v NP-40, 10 mM Tris HCl, pH 8.0); and 1X TE buffer once. Beads were resuspended in 240 µL TE + 1% SDS and incubated at 65°C for 10 min with frequent vortexing. Eluates were transferred to new tubes. Beads were again resuspended in 240 µL TE + 1% SDS and incubated at 65°C for 10 min with frequent vortexing. The second eluate was combined with the first and brought up to a total volume of 800 µL. Input samples were also brought up to a total volume of 800 µL. IP and input samples were incubated with 40 µL RNase A (Invitrogen #AM2271) at 37°C for 30 min. 40 µL Proteinase K (Invitrogen #AM2548) was added, and samples were incubated at 65°C overnight to reverse crosslinks. DNA was harvested and purified with the ChIP DNA Clean & Concentrator kit (Zymo Research #D5205) according to the manufacturer’s instructions and eluted in 22 µL elution buffer. 1 µL was used for quantification with a 1X dsDNA High Sensitivity kit (Invitrogen #Q33231) on a Qubit Flex Fluorometer device (Invitrogen #Q33327), and the rest was submitted for sequencing through Azenta Life Sciences.

### ChIP-seq Analysis

Library preparation and sequencing were performed by Azenta Life Sciences. ChIP DNA samples were quantified using a Qubit 3.0 Fluorometer (Invitrogen #Q33216) and DNA integrity was verified with the 4200 TapeStation System (Agilent #G2991BA). The NEBNext Ultra II DNA Library Prep kit for Illumina (New England Biolabs #E7645) was used following the manufacturer’s recommendations. Briefly, ChIP DNA was end-repaired, and adapters were ligated after adenylation of the 3’ ends. Adapter-ligated DNA was size selected, followed by cleanup and limited cycle PCR enrichment. The ChIP library was validated using the TapeStation System and quantified using a Qubit 3.0 Fluorometer and real-time PCR. The sequencing libraries were multiplexed and clustered onto a flowcell on a NovaSeq X Plus instrument (Illumina) and sequenced in a 2×150-bp configuration, generating 150-bp paired-end reads at a depth of 16–24 million reads per sample. Image analysis and base calling were conducted by NovaSeq Control Software (NCS). Raw sequence data (.bcl files) were converted into FASTQ files and demultiplexed using Illumina bcl2fastq 2.20 software. One mismatch was allowed for index sequence identification. The BBMap package (v38.18) ^83^ was used to remove duplicates and trim adapters from the reads with the clumpify and BBDuk tools respectively. Quality trimming was performed with Trimmomatic (v0.39) ^84^. Sequence reads were aligned to the human reference genome (hg38) with BWA-MEM (v0.7.17) ^85^. Aligned reads were filtered, using a Mapping Quality (MAPQ) score of at least 20, sorted, and indexed with SAMtools ^86^. Peak calling was performed for each replicate and merged alignments using MACS2 ^87^ with default settings. Differential binding analysis comparing locus-specific versus sham gRNAs was performed on consensus peaks using DiffBind ^88^ with significance thresholds of adjusted p value (FDR) <0.01 and fold change >2. To control for batch effects, the design for the differential binding analysis was defined as “∼ Factor + Condition”, where Factor is the batch and Condition is either the locus-specific or sham gRNA, with the dba.contrast function. Consensus peaks and differentially bound peaks were annotated for genomic features using ChIPpeakAnno ^89^. For visualization of genomic tracks, the aligned reads were converted to bedGraph files and normalized by reads per million (RPM) with BEDTools ^90^. BedGraph files were converted to bigWig format using the bedGraphToBigWig utility provided by the UCSC Genome Browser ^91^. ChIP-seq tracks were visualized using Integrative Genomics Viewer (IGV) version 2.19.7 ^92^. A complete list of peaks identified in the differential binding analysis is provided in **Supplemental Table 2**.

### Immunofluorescence

Cells were plated on glass coverslips inside 6-well cell culture plates. Cells were fixed in 4% paraformaldehyde (Thermo Fisher #J19943.K2) with gentle shaking for 30 min followed by 100% methanol at -20°C for 10 min. Coverslips were blocked in a solution of 10% milk in PBSTx buffer at room temperature for 1 hour, then incubated overnight at 4°C with primary antibodies, followed by 1 hour at room temperature with secondary antibodies (listed in **Supplemental Table 1**) and DAPI stain (Sigma-Aldrich #D9542). Coverslips were mounted on slides with ProLong Diamond Antifade (Invitrogen #P36961). Slides were imaged using a 40X oil objective on a Leica SP8 laser microscope equipped with HyD detectors individually tuned to each fluorophore. Channels were acquired separately to minimize crosstalk.

### C-BERST Biotinylation Assay

C-BERST was performed as previously described by Gao et al ^28^ with the following modifications. Following 21 hours of doxycycline and Shield1 induction (see *Plasmids*), 60 million HT29 (WT APC) cells expressing dCas9-APEX2 and gRNAs were incubated with 500 µM biotin phenol (BP) (Sigma-Aldrich #SML2135) for 30 min at 37°C. 1 mM H_2_O_2_ (Sigma-Aldrich #H1009) was added, and cells were placed for 1 minute on a horizontal shaker at room temperature. Cells were washed 3 times with antioxidant quencher solution (5 mM Trolox, Thermo Scientific Chemicals #218940050; 10 mM sodium ascorbate, Thermo Scientific Chemicals #AAA1775922; 10 mM sodium azide, Thermo Scientific Chemicals #AC190381000), and 2 times with PBS. Cells were scraped in 15 mL PBS and centrifuged at 300g for 5 min. Pellets were washed in 10 mL PBS and centrifuged again at 300g for 5 min. Pellets were resuspended in 2 mL Nuclei EZ Lysis Buffer (Sigma-Aldrich #NUC101) containing 1X Halt Protease and Phosphatase Inhibitor Cocktail, vortexed, incubated on ice for 5 min, then centrifuged at 300g for 5 min at 4°C. Pellets were again resuspended in 2 mL Nuclei EZ Lysis Buffer with 1X protease inhibitor, vortexed, incubated on ice for 5 min, then centrifuged at 300g for 5 min at 4°C. Nuclear pellets were resuspended in 500 µL RIPA lysis buffer (50 mM Tris HCl, 150 mM NaCl, 0.125% SDS, 0.125% sodium deoxycholate, 1% Triton X-100, pH 7.5) with 1X protease inhibitor and vortexed well. Nuclei were sheared by pipetting through an insulin syringe, incubating on ice for 10 min, and repeating for a total of 3 cycles. Lysates were sonicated for 15 min using a Q800R2 sonicator (QSonica) with 20 seconds on/20 seconds off cycling. Lysates were incubated overnight on a tube rotator at 4°C with 450 µL Dynabeads MyOne Streptavidin T1 (Invitrogen #65602) pre-equilibrated in 3 washes of 0.2% BSA in PBS. The following day, beads were washed with the following buffers: 2 times with RIPA lysis buffer; 1 time with 1 M KCl; 1 time with 0.1 M Na_2_CO_3_; and 2 times with 2 M urea in 10 mM Tris HCl, pH 8.0. Wash solution was completely removed, and dried beads were stored at -80°C, then submitted for LC-MS/MS through the Wistar Institute Proteomics and Metabolomics Facility.

### Protein Mass Spectrometry

Two independently cultured biological replicates per condition were submitted for mass spectrometry. Bead-bound samples were washed, reduced with TCEP, alkylated with iodoacetamide, and digested off the beads using trypsin. Digests were desalted on C18 spin columns, then analyzed using a 120-minute LC-MS/MS (OT-IT) data-dependent acquisition method on an Orbitrap Eclipse mass spectrometer (Thermo Scientific) with FAIMS. Data analysis was performed using Proteome Discoverer v3.1.1.93 (Thermo Scientific). Spectra were searched against the Swiss-Prot Human database (UniProt consortium) with curated isoforms (UP000005640, downloaded 08-06-2024), a custom database containing the recombinant sequences of the WT APC and dCas9-APEX2 constructs, and a common contaminant database using Sequest HT. Common contaminants were removed, as were proteins identified by a single unique peptide. Final data were reported as abundance values normalized to the total protein abundance of each sample. Comparisons were performed on log2-transformed normalized abundance values between groups to report a fold change and moderated t-test p value adjusted for multiple testing using Benjamini-Hochberg FDR correction (p_adj_). Proteins with significant changes between groups satisfied the criteria of (1) at least a twofold change in abundance, and (2) p_adj_<0.05. Proteins with no reported nuclear localization were included in all statistical analyses but are not depicted. A complete list of proteins identified in the analysis is provided in **Supplemental Table 3**.

### Luciferase Reporter Assay

LS174T (DNTCF) cells were plated in white-bottom 96-well plates. 100 ng of *Renilla* luciferase and 200 ng of firefly luciferase plasmids were co-transfected (see *Plasmids*). Following DNTCF induction by DOX, cells were lysed and luminescence was read using the Dual-Glo Luciferase Assay System (Promega #E2940) on a FLUOstar Omega plate reader (BMG Labtech). Firefly readings were normalized to *Renilla* activity from the same well at the assay endpoint.

### siRNA Screen

A custom array of Silencer Select siRNAs consisting of 2 siRNAs per target was obtained (Invitrogen). siRNAs used in the screen are listed in **Supplemental Table 1**. 5 nM siRNA was transfected into LS174T (DNTCF) cells with Lipofectamine RNAiMAX Transfection Reagent (Invitrogen #13778075) according to the manufacturer’s instructions. Cells were harvested for analysis 72 hours following transfection.

### CRISPR/Cas9

p38α knockout LS174T (DNTCF) and HT29 (WT APC) cells were generated previously using Cas9 and gRNA RNP electroporation with the 4D-Nucleofector System (Lonza Bioscience) ^20^. Cells electroporated with Cas9, but without gRNA, were used as wild-type controls.

### Colonic Organoid Culture

Animal protocols were approved by the Thomas Jefferson University Institutional Animal Care and Use Committee (IACUC protocol number 00914-2). C57BL/6(J) mice (Jackson Laboratory #000664) were euthanized at 6–12 weeks of age. The proximal colon was collected, flushed with cold PBS, minced, and placed into a 10 mL solution of Advanced DMEM/F12 (Gibco #12634028), 1% Penicillin-Streptomycin (Gibco #15140122), 0.5% Gentamicin (Gibco #15750060), 1% FBS, and 15 mM EDTA. Tissue was placed on a tube rotator at 4°C for 1 hour. Tissue was vortexed twice for 1 minute each to isolate colonic crypts. Large debris was removed and crypts were centrifuged at 500g for 3 min at 4°C. Crypts were washed twice in a 5 mL solution of Advanced DMEM/F12, 1% Penicillin-Streptomycin, 0.5% Gentamicin, 1% FBS, and 2% sorbitol (Sigma-Aldrich #S1876). Crypts were washed once in Advanced DMEM/F12, then resuspended in Matrigel Growth Factor Reduced Basement Membrane Matrix (Corning #356231). 50 μL Matrigel/crypt droplets were plated in 24-well plates and cultured with 500 μL of medium consisting of 50% conditioned medium from L-WRN cells (ATCC #CRL-3276, cultured and conditioned according to manufacturer instructions) supplemented with 1X Antibiotic-Antimycotic (Gibco #15240112), 10 mM HEPES, 1% GlutaMAX (Gibco #35050061), 1X N-2 (Gibco #17502001), 1X B-27 (Gibco #17504001), 0.1 mM N-acetyl-L-Cysteine (Cayman Chemical #20261), 0.05 μg/mL EGF (Gibco #AF-100-15), 0.5 μM A 83-01 (StemCell Technologies #72022), 10 nM Gastrin-1 (AnaSpec #AS-64149), 10 μM Y-27632 (StemCell Technologies #72304), and 10 μM CHIR99021 (StemCell Technologies #72054). Medium was replaced every 3 days and organoids were passaged weekly. For differentiation studies, differentiation medium was used for 3 days, consisting of Advanced DMEM/F12 supplemented with the above reagents, in addition to 0.1 μg/mL Noggin (Gibco #120-10C) and 1 μg/mL R-Spondin 1, (Gibco #120-38), and without Gastrin, Y-27632, or CHIR99021. Organoids were harvested for protein analysis by incubating Matrigel patties in 300 μL Cell Recovery Solution (Corning #354253) for 30 min at 4°C, centrifuging at 200g for 5 min, and resuspending in M-PER buffer as described above.

### Co-immunoprecipitation (co-IP)

Co-immunoprecipitation was performed in LS174T (DNTCF) cells expressing pFLAG-GATA2 (see *Plasmids*) using the Nuclear Complex Co-IP Kit (Active Motif #54001) according to the manufacturer’s instructions. Briefly, ∼8.8 million cells were lysed, and nuclei were isolated as described. 500 μg of nuclear lysate was incubated with 5 μg antibody (**Supplemental Table 1**) overnight at 4°C on a tube rotator. The following day, the complex was incubated on Protein G Agarose Columns (Active Motif #53039) for one hour at 4°C on a tube rotator. Low-stringency buffer supplemented with 150 mM NaCl and 1% detergent was used for washes. Protein was eluted from columns using 2X Reducing Buffer. Nuclear inputs and IP samples were boiled at 95°C for 5 min and loaded onto gels for immunoblot as described above.

### Statistics and Data Availability

Results are provided as mean ± standard deviation (SD), and a p value of <0.05, or <0.01 where indicated, was considered significant. Statistical analysis was performed in GraphPad Prism version 11.0.1 using analyses indicated in each figure legend, including Welch’s t-tests and two-way ANOVA with corrections for multiple comparisons. A complete record of proteomics and ChIP-sequencing data are provided in supplementary materials. Additionally, ChIP-sequencing data have been deposited to the Gene Expression Omnibus (GSE348607) and will be made publicly available upon publication of the manuscript.

## Supplemental Figures & Legends

**Supplemental Figure 1:**
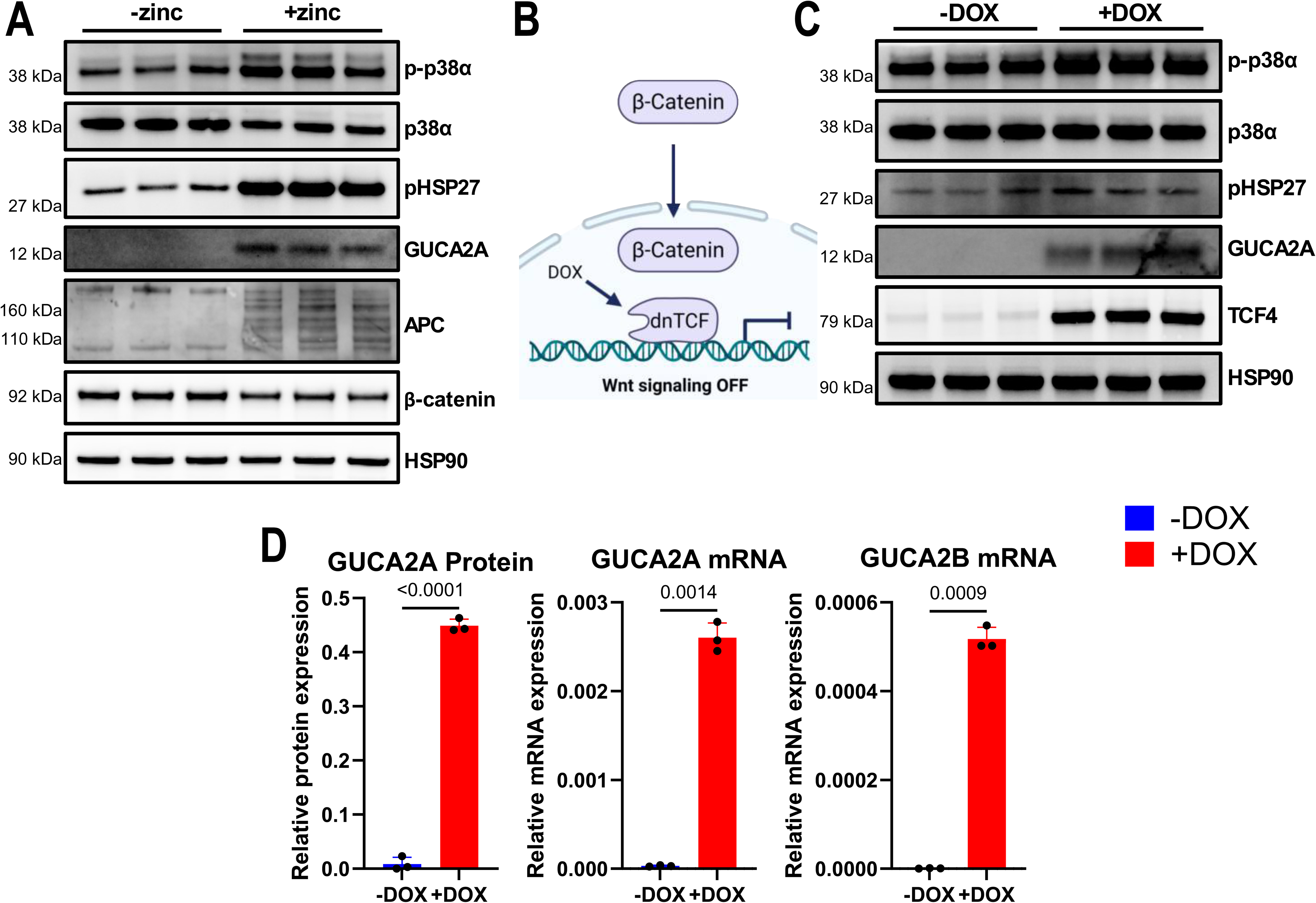
GUCA2A and GUCA2B induction by WT APC and DNTCF. (A) Full immunoblot of HT29 (WT APC) cells with and without 24 hours of zinc treatment. (B) Schematic of LS174T (DNTCF) cells. Following 48 hours of doxycycline (DOX) treatment, dominant negative TCF4 (DNTCF) is induced, preventing β-catenin binding and suppressing downstream β-catenin/TCF signaling. (C) Full immunoblot of LS174T (DNTCF) cells with and without 48 hours of DOX treatment. (D) GUCA2A protein and mRNA, and GUCA2B mRNA, are induced following DOX induction of DNTCF. Welch’s t-test. N=3 biological replicates per condition (A, C-D) with 2 technical replicates per sample in qPCR (D).

**Supplemental Figure 2:**
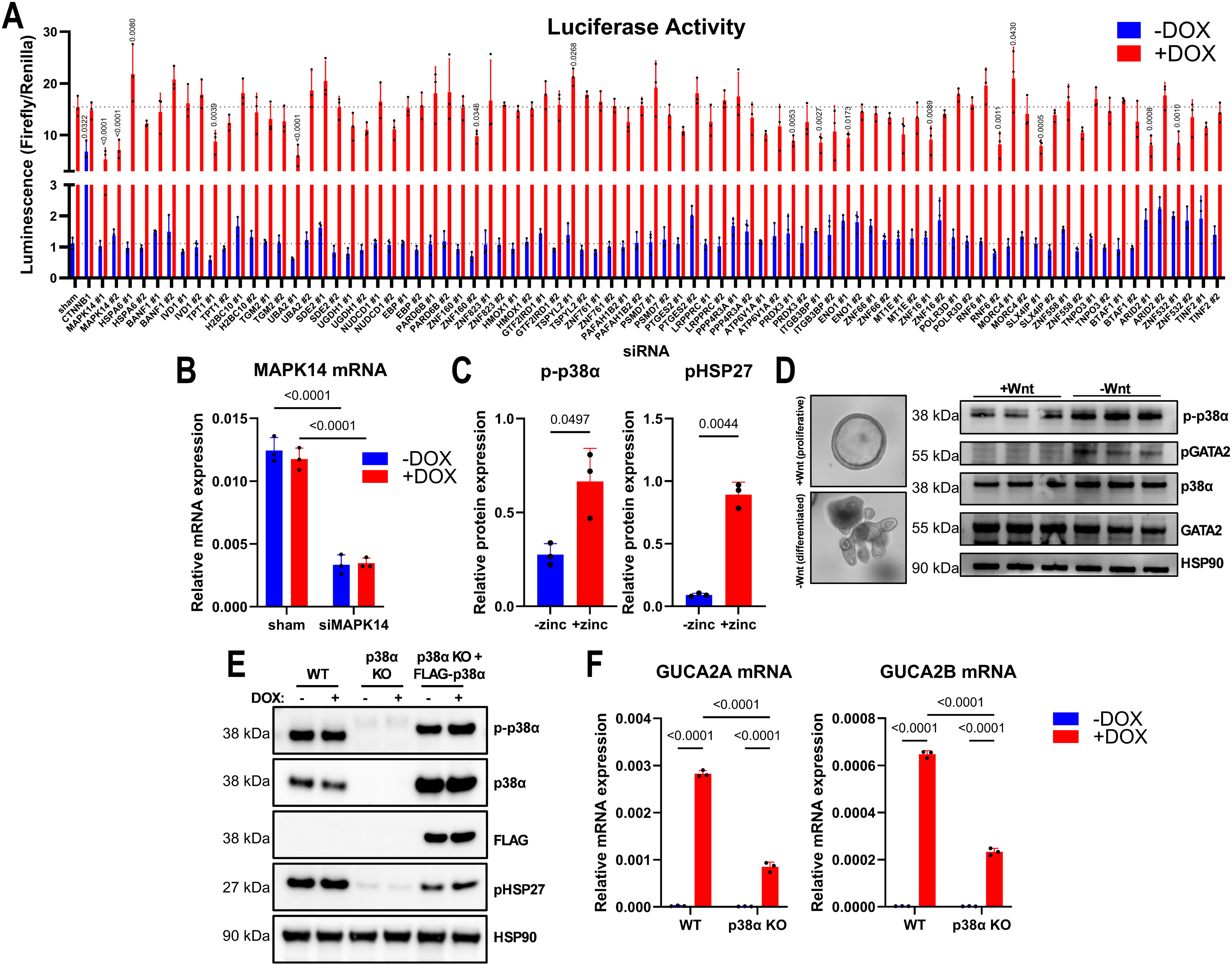
Luciferase screen reveals p38α as a regulator of GUCA2A and GUCA2B. (A) Luciferase activity in LS174T (DNTCF) cells of a 6-kb *GUCA2A* reporter vector with a screen of siRNAs against candidates from LC-MS/MS. Two siRNAs were used individually per candidate, and a significant effect by both siRNAs was required to prioritize candidates. β-catenin (CTNNB1) was used as a control for a negative regulator of GUCA2A. Shown p values compare each siRNA to the sham siRNA under the same condition (with or without DOX). Complete list of candidates and their directionality can be found in Supplemental Table 1. (B) MAPK14 mRNA expression is suppressed by MAPK14 siRNA (siRNA ID s3586) in LS174T (DNTCF) cells. (C) Phosphorylated p38α (T180/Y182) and downstream HSP27 (S82) are suppressed by β-catenin/TCF signaling in HT29 (WT APC) cells. Quantified from immunoblot in Supplemental Figure 1A. (D) Immunoblot of mouse colonic organoids cultured in Wnt-containing proliferative medium or Wnt-free differentiation medium. (E) Immunoblot of LS174T (DNTCF) wild-type (WT), p38α knockout (KO), and KO cells expressing FLAG-p38α. (F) GUCA2A and GUCA2B expression is reduced by p38α knockout in LS174T (DNTCF) cells. Two-way ANOVA with Dunnett’s correction for multiple comparisons (A); two-way ANOVA with Tukey’s correction for multiple comparisons (B, F); Welch’s t-test (C). N=3 biological replicates per condition (A-F) with 2 technical replicates per sample in qPCR (B, F).

**Supplemental Figure 3:**
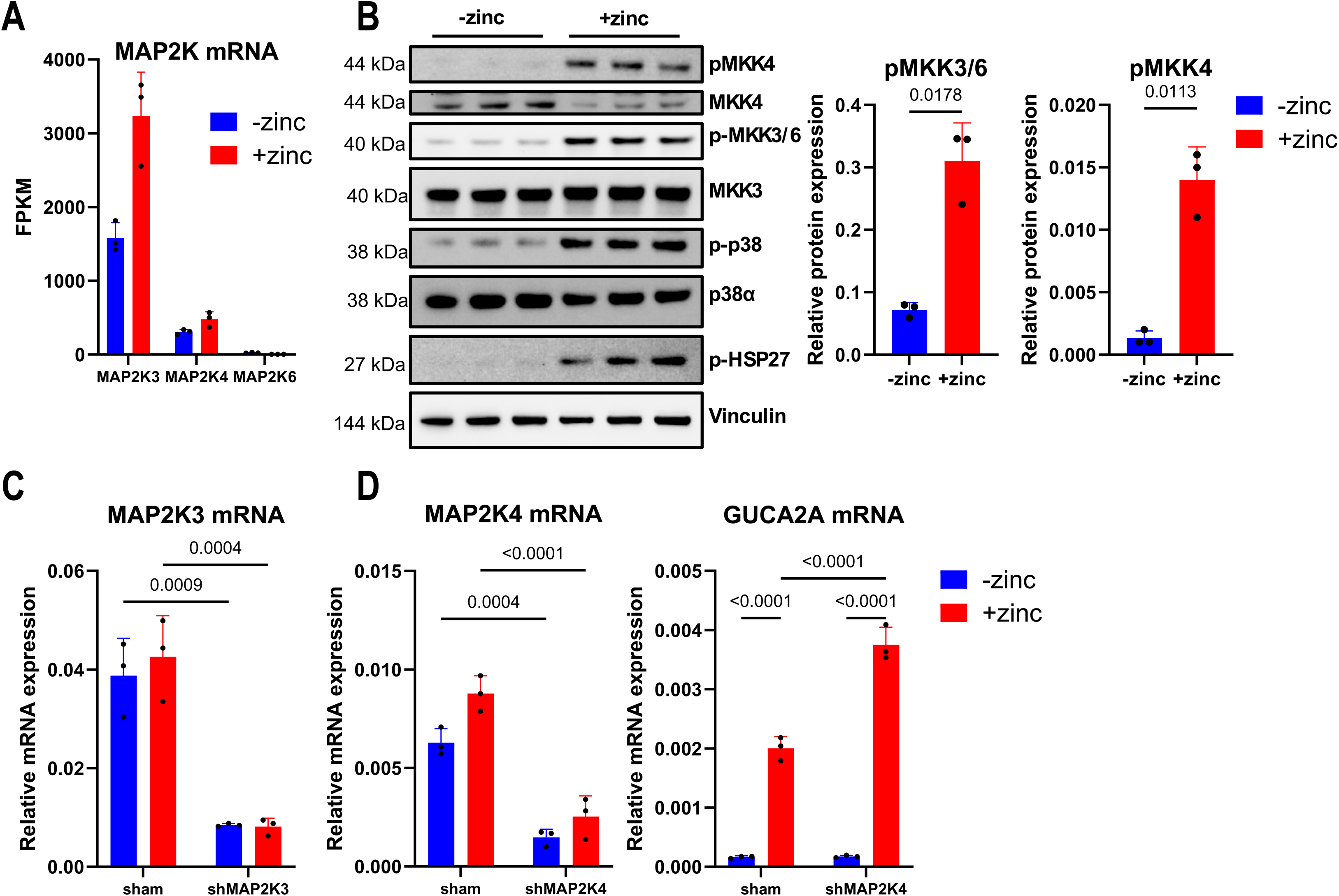
p38α upstream activity is impaired by β-catenin/TCF signaling. (A) Expression of MKK3 (MAP2K3), MKK4 (MAP2K4), and MKK6 (MAP2K6) in HT29 (WT APC) cells from RNA-seq ^18^. Reads in FPKM. (B) Immunoblot (left) and quantification (right) of phosphorylated MKK3/6 and phosphorylated MKK4 in HT29 (WT APC) cells. (C) MAP2K3 mRNA expression is suppressed by MAP2K3 shRNA in HT29 (WT APC) cells. (D) MAP2K4 (left) and GUCA2A (right) mRNA in the presence of MAP2K4 shRNA. Welch’s t-test (B); two-way ANOVA with Tukey’s correction for multiple comparisons (C-D). N=3 biological replicates per condition (A-D) with 2 technical replicates per sample in qPCR (C-D).

**Supplemental Figure 4:**
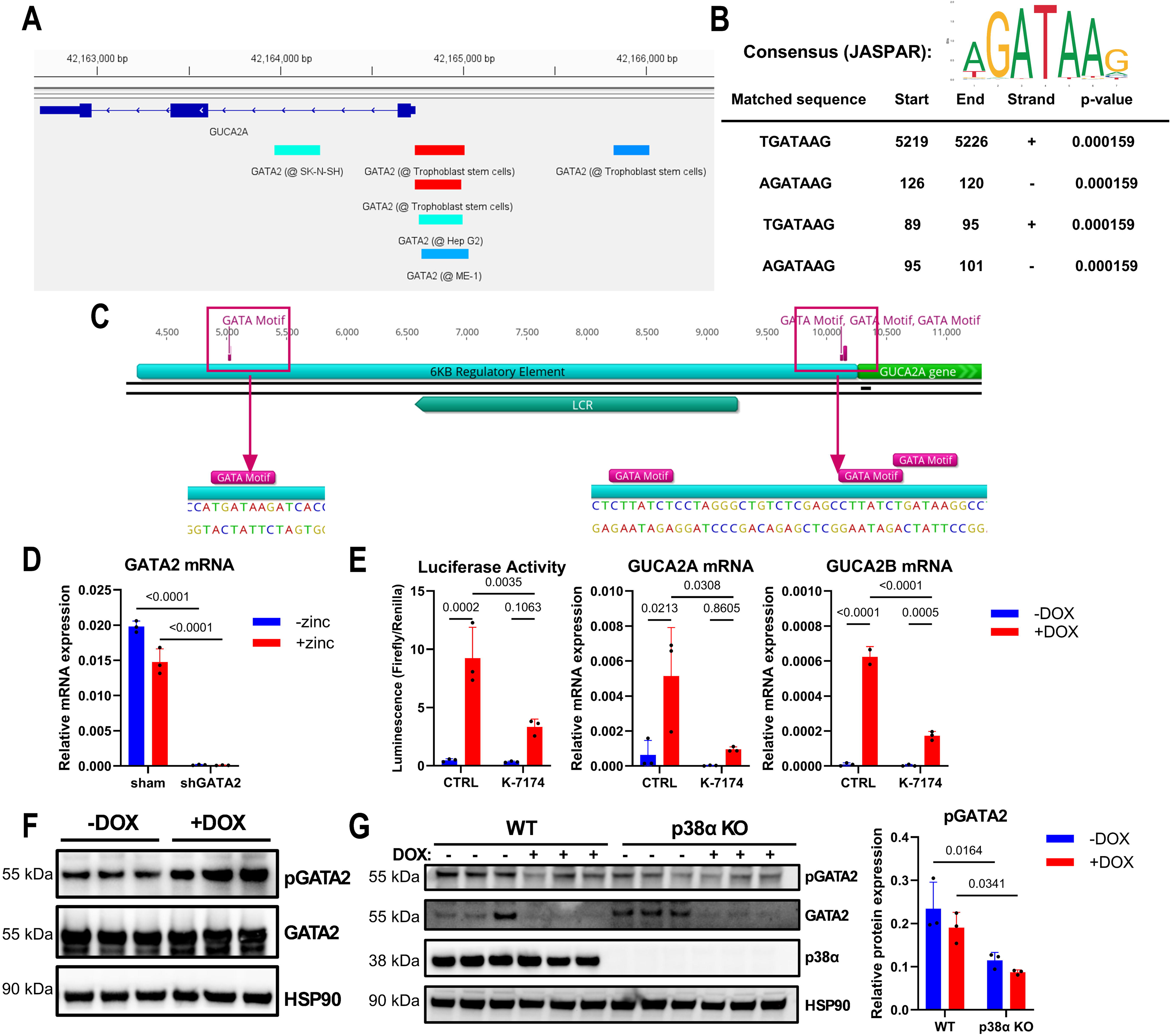
GATA2 regulates GUCA2A and GUCA2B. (A) Snapshot of anti-GATA2 ChIP-seq peaks at the *GUCA2A* locus from the ChIP-Atlas. Models used in the assays are denoted. Peak color represents Q value calculated by MACS2 peak-caller; blue (<1E-05), cyan (<1E-25), red (<1E-100) ^70^ ^87^. (B) FIMO (Find Individual Motif Occurrences) motif search of the GATA2 consensus motif from JASPAR. Listed start sites are upstream from the *GUCA2A* transcription start site. (C) Snapshot of the identified GATA2 consensus motifs in (B) relative to the *GUCA2A* locus. The entire 6-kb regulatory element as well as the LCR are indicated. Insets show a zoomed-in view of the four motif occurrences. (D) GATA2 mRNA expression is suppressed by GATA2 shRNA in HT29 (WT APC) cells. (E) GUCA2A luciferase reporter activity, GUCA2A mRNA, and GUCA2B mRNA are suppressed by treatment with 20 µM of the GATA inhibitor K-7174 in LS174T (DNTCF) cells. (F) Immunoblot of LS174T (DNTCF) cells expressing FLAG-tagged GATA2. (G) Native GATA2 immunoblot in wild-type and p38α KO LS174T (DNTCF) cells. Quantification of phosphorylated GATA2. Two-way ANOVA with Tukey’s correction for multiple comparisons (D-E, G). N=3 biological replicates per condition (D-G) with 2 technical replicates per sample in qPCR (D-E).

## Supplemental Tables

**Table S1:** Materials and reagents.

**Table S2:** Differential binding peaks identified by dCas9-APEX2 ChIP-seq.

**Table S3:** C-BERST proteomics results.

## Acknowledgments

HT29 (WT APC) cells were provided by B. Vogelstein ^33^. LS174T (DNTCF) cells were provided by H. Clevers ^54^. pEJS578_DD-dSpyCas9-mCherry-APEX2 was a gift from E. Sontheimer (Addgene #108570; http://n2t.net/addgene:108570; RRID:Addgene_108570). pCDNA3 Flag p38 alpha was a gift from R. Davis (Addgene #20351; http://n2t.net/addgene:20351; RRID:Addgene_20351). pFLAG-GATA2 was a gift from G. Hotamisligil (Addgene #1418; http://n2t.net/addgene:1418; RRID:Addgene_1418). ChIP-seq was performed by Azenta Life Sciences. LC-MS/MS and proteomic analysis were performed by the Wistar Institute Proteomics and Metabolomics Facility. ChIP-seq tracks were visualized using Integrative Genomics Viewer (IGV) version 2.19.7 ^92^. Publicly available ChIP-seq data were accessed using ChIP-Atlas ^70^. Transcription factor binding motifs were obtained from the JASPAR 2026 database ^71^ and interrogated using FIMO motif analysis through the MEME Suite ^72^. Genomic and plasmid DNA sequences were visualized on Geneious Prime 2026.0. Protein phosphorylation data were obtained from the PhosphoSitePlus database ^77^. Figures and statistical analyses were generated using GraphPad Prism version 11.0.1. Schematics were created with BioRender.com. The authors would like to thank A. Faustino and H.Y. Tang for their expertise and assistance with LC-MS/MS and proteomic analysis; the SKCCC Division of Biostatistics and Bioinformatics for performing ChIP-seq and RNA-seq data analysis and for their expertise in computational biology; and T. Baybutt for providing expertise in plasmid cloning and lentivirus production.

